# CELF2 suppresses endogenous RNA ligands that activate RIG-I-mediated interferon induction

**DOI:** 10.1101/2025.03.27.645787

**Authors:** Julian R. Smith, Olivia C. Kern, Gargi Mishra, Nandan S. Gokhale, Noa Etzyon, Kim Somfleth, Johannes Schwerk, Kenji Nakamichi, Cole M. Pugliano, Zachary Lindbloom-Brown, Michael Gale, Ram Savan

## Abstract

Type I interferons (IFNs) are critical for the control of viral infections, but aberrant IFN expression can result in tissue damage. RIG-I-like receptors (RLRs), such as RIG-I and MDA5, sense viral RNA and signal through the adaptor protein MAVS to induce phosphorylation and nuclear translocation of the transcription factor IRF3, thereby driving IFN production. However, activation of RLRs by endogenous RNA ligands can also induce IFNs, leading to autoinflammation. Identifying factors that suppress endogenous RNA ligands is critical for preventing IFN-induced autoimmunity. In this study, we identify a novel regulator, CELF2, that suppresses endogenous RNAs that otherwise activate the RLR pathway. We uncovered a novel role for the splicing factor CELF2 as a suppressor of immunostimulatory endogenous RNA ligands. Depletion of CELF2 in macrophages led to a spontaneous IFN and IFN-stimulated gene signature, dependent on the RIG-I-MAVS pathway. Furthermore, the transfer of RNA from CELF2-depleted macrophages was sufficient to induce type I IFN expression in naïve cells. This RNA was found to be double-stranded as RNase III treatment of RNA derived from CELF2-depleted cells ablated IFN induction in naïve cells. Immunoprecipitation of double-stranded RNA from CELF2-depleted macrophages revealed several immunostimulatory RNAs, which contribute to the increased interferon-stimulated gene signature observed in CELF2-depleted macrophages. These data indicate that CELF2 suppresses endogenous RNA ligands, which could otherwise activate RIG-I and induce an IFN signature. Overall, these findings reveal that CELF2 is an important regulator of self-RNA ligands to prevent IFN-induced autoinflammation.

**One-sentence summary:** CELF2 suppresses RIG-I-like receptor ligands that activate interferon.

## Introduction

While type I interferons (IFNs) are critical for controlling viral infection, unchecked IFN signaling can lead to interferonopathies(*1-3*). There is strong evidence that endogenous RNA ligands can activate interferon through RIG-I-like receptors (RLRs: RIG-I and MDA5)(*4*). Endogenous RNA ligands that activate RLRs have been discovered, some within the context of viral infections (reviewed in *5*). Viruses can promote the unmasking or mis-localization of ribosomal RNA pseudogenes and vault RNA from ribonuclear proteins or lead to the recognition of 5’-triphosphate (5’-ppp) and poly-uridine motifs(*6*) recognized by RIG-I and promote antiviral signaling and viral clearance(*7, 8*). Sensing of endogenous RNA ligands containing Alu-rich repeats has also been implicated in IFN-mediated diseases. While the specific identities of these Alu-rich ligands have not been well described, studies investigating loss-of-function ADAR1 mutations and gain-of-function MDA5 mutations have identified increased sensing of endogenous RNAs with Alu element repeats(*9-13*). Loss of function mutations in SKIV2L and PNPase, proteins involved in RNA processing and the metabolism of endogenous immunostimulatory RNA ligands, result in IFN production and damaging inflammation(*14-16*). Together, these findings support a hypothesis that endogenous RNA can act as damage-associated molecular patterns (DAMPs) and, depending on the context, induce interferons driven by RLRs. However, our understanding of how immunostimulatory endogenous RNAs are generated and the host factors that suppress their biogenesis are unclear.

RNA splicing factors play important roles in post-transcriptional programming of immune responses(*17-19*). Indeed, viruses often co-opt or disrupt splicing factor function to support their own replication, positioning these factors as key regulators of the inflammatory response(*20-23*). In addition, chemical inhibition of splicing or depletion of splicing factors can generate dsRNA ligands that result in IFN activation(*24-26*). In this study, we identify a novel role for the splicing factor CELF2 in the suppression of dsRNA that leads to spontaneous IFN production in macrophages. CELF2 is a member of the CUG-BP Elav-like family of RNA-binding proteins, and is a regulator of RNA processing, including pre-mRNA alternative splicing, alternative polyadenylation(*27*) and mRNA decay(*28, 29*). Here we found that the absence of CELF2 generates endogenous-double-stranded RNA which is sufficient to activate RLR responses. We observed increased RNA:RIG-I interactions in the absence of CELF2 as well as IRF3 activation and downstream IFN signaling. RNA sequencing analysis of dsRNA in CELF2-depleted cells found several unique dsRNAs enriched. Depletion of several of these dsRNAs reduced the interferon signature found in CELF2-depleted macrophages. These data provide a novel role for CELF2 in suppressing IFN activation in macrophages by suppressing immunostimulatory endogenous RNA.

## Results

### Loss of CELF2 results in spontaneous production of type I IFN and an ISG signature

To investigate the contributions of CELF2 to innate immunity and splicing, we used siRNA to transiently deplete CELF2 in U-937 macrophages differentiated from monocytes by Phorbol 12-myristate 13-acetate (PMA) stimulation (Fig. 1A). For the rest of the manuscript, we will describe PMA-differentiated U-937 cells as macrophages. We transiently depleted CELF2 as *Celf2*^-/-^ mice have developmental defects and do not survive past embryonic stages, as is common following the deletion of other splicing factors(*30, 31*). We performed long-read RNA sequencing to assess differential gene expression and alternative isoform usage upon CELF2 depletion. We found that depletion of CELF2, in the absence of any innate immune stimulus, resulted in robust type I IFN induction and an associated interferon-stimulated gene (ISG) signature (Fig. 1B-C). These findings implicated CELF2 as a negative regulator of spontaneous IFN production.

**Figure 1.**
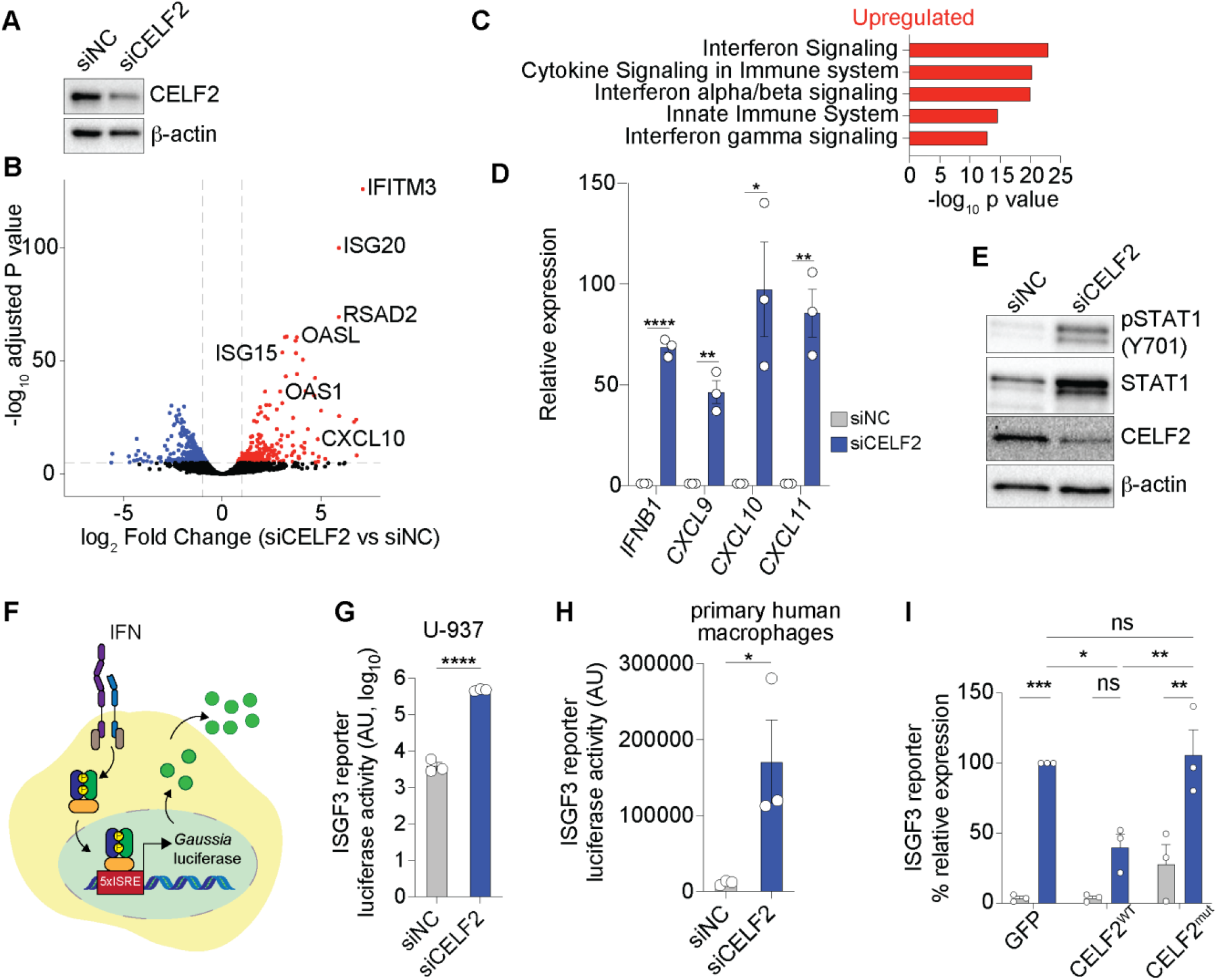
CELF2-depleted macrophages exhibit a spontaneous interferon-stimulated gene signature. A. Representative immunoblot analysis of CELF2-depleted (siCELF2) and control (siNon-targeting Control (NC)) macrophages. B. Volcano plot of differentially expressed genes in CELF2-depleted macrophages compared to control macrophages from Oxford Nanopore Technologies MinION long-read RNA sequencing. Red indicates significantly upregulated genes; blue indicates significantly downregulated genes. p < 0.05, n=3. C. Reactome pathway analysis of significantly upregulated genes from long-read RNA sequencing analysis. D. RT-qPCR analysis of *IFNB1, CXCL9, CXCL10* and *CXCL11* induction following CELF2 depletion relative to housekeeping gene *HPRT1*. E. Immunoblot analysis of STAT1 phosphorylation following CELF2 depletion. F. Schematic of 5xISRE *Gaussia* luciferase reporter system. IFN stimulates the production of *Gaussia* Luciferase downstream of 5xISRE reporter that is specifically induced by ISGF3. *Gaussia* luciferase can be measured in the supernatant. G. 5xISGF3 *Gaussia* Luciferase reporter activity of supernatants from CELF2-depleted U-937 cells. Supernatants were taken from CELF2-depleted U-937 macrophages and transferred onto ISGF3 luciferase reporter cells and luciferase activity was measured after 24h. H. 5xISGF3 *Gaussia* luciferase reporter activity from CELF2-depleted primary human macrophage supernatants. I. 5xISGF3 *Gaussia* luciferase reporter activity from CELF2-depleted cells engineered to express GFP, CELF2^WT^ or CELF2 RNA-binding mutants (CELF2^mut^). Data in all panels are representative of n=3 independent experiments. Each symbol represents an individual biological replicate. Graphs display mean ± SEM. Student’s t-test (D, G, and H), Two-way ANOVA (I); * represents p < 0.05, ** represents p ≤ 0.01 **** represents p ≤ 0.0001, ns represents p > 0.05. A and E immunoblots representative of n=3 independent experiments.

To confirm our long-read RNA-seq results, we measured significant increases in *IFNB1, and* IFN induced (*32*) *CXCL9, CXCL10* and *CXCL11* mRNA abundance by RT-qPCR (Fig. 1D) and STAT1 phosphorylation by immunoblot (Fig. 1E) following CELF2 depletion. These findings were also confirmed using an siRNA pool from a different source which targeted different sites within the *CELF2* transcript (Fig. S1B). In addition, we confirmed CELF2-dependent IFN and ISG induction in another macrophage-like cell line, THP-1 (Fig. S1C-F). To confirm that IFN was being secreted, we transferred supernatants from U-937 cells onto a reporter cell line which secretes *Gaussia* luciferase downstream of a 5xISRE promoter in response to IFN signaling (Fig. 1F)(*33*). We found that supernatants from CELF2-depleted cells resulted in a robust increase in luciferase activity, demonstrating that IFN is secreted from CELF2-deficient macrophages (Fig. 1G). Furthermore, we depleted CELF2 using a *CELF2*-targeting siRNA in primary M-CSF-differentiated monocytes and found that CELF2 depletion in primary macrophages resulted in IFN secretion (Fig. 1H). Together, these data suggest that loss of CELF2 in macrophages results in the spontaneous production of IFN.

The expression of most CELF proteins is tissue-restricted following development(*34*). CELF2 is robustly expressed in immune cells, cells of the central nervous system, the developing heart, and in stem cells(*35*). To rule out off-target effects of the *CELF2*-targeting siRNA, we used Huh7 hepatocytes which do not express CELF2 (Fig. S1G). Introducing *CELF2*-targeting siRNA into Huh7 cells did not increase *IFNB1* or *CXCL10* expression or STAT1 phosphorylation (Fig. S1G-I). We next tested if the RNA-binding function of CELF2 was required to suppress the spontaneous production of IFN by complementation of wild type CELF2 (CELF2^WT^) or CELF2 RNA-binding mutant (CELF2^mut^) in CELF2-depleted cells. CELF proteins bind RNA *via* three conserved RNA recognition motifs (RRMs) and enable splicing, alternative polyadenylation, or regulate mRNA translation(*28, 29*). We generated RNA-binding mutant CELF2 by mutating five key residues in the RNA recognition motifs (RRMs) that have previously been shown to be important for RNA binding in the closely related CELF1 protein(*36, 37*) (Fig. S1J). To confirm that the RNA-binding CELF2 mutants lack the ability to bind RNA, we overexpressed FLAG-tagged CELF2^WT^ or CELF2^mut^ in HEK293T cells, immunoprecipitated these proteins and isolated bound RNA (Fig. S1K). Immunoprecipitation of CELF2^WT^ enriched a representative transcript *PQBP1* compared to the empty vector (EV) control (Fig. S1L). However, CELF2^mut^ failed to pulldown *PQBP1* transcript, similar to the EV control (Fig. S1L), demonstrating that the 5 mutations in the RRMs were sufficient to prevent CELF2-RNA binding. We then transduced U-937 cells to stably express CELF2^WT^ or CELF2^mut^, as well as GFP as a control, and depleted CELF2 as previously described. As expected, depletion of CELF2 in GFP expressing control cells resulted in IFN production as measured by supernatant transfer onto 5xISGF3-GLuc reporter cells (Fig. 1I). However, supernatants from CELF2-depleted cells which constitutively express CELF2^WT^ demonstrated a significant reduction in IFN reporter activity compared to control GFP cells and no difference compared to non-targeting siRNA control treated cells (Fig. 1I). Depletion of CELF2 in CELF2^mut^ expressing cells did not suppress IFN reporter activity indicating that CELF2 RNA-binding ability is required to suppress IFN production in macrophages (Fig. 1I). Together these data demonstrate that RNA-binding ability of CELF2 is required for the suppression of type I IFN in macrophages.

### Spontaneous IFN production upon *CELF2*-depletion is MAVS dependent

IFN production in macrophages can be triggered by nucleic acid sensing through the RIG-I-like receptor (RLR)-MAVS or the cGAS-STING pathway. These two pathways converge on the activation of IRF3, the master transcription factor for IFN induction(*38, 39*). We found that depletion of CELF2 induced phosphorylation of IRF3 and STAT1 proteins (Fig. 2A). To test if IRF3 was required for the spontaneous induction of IFN following CELF2 depletion, we generated IRF3 knockout (KO) U-937 cells using CRISPR-Cas9 technology and depleted CELF2 in these cells. As expected, non-targeting control cells exhibited a significant increase in *IFNB1* transcription and STAT1 phosphorylation following CELF2 depletion, but these increases were ablated in IRF3 KO cells (Fig. 2B-C). These data suggested that IFN production induced by CELF2 depletion is dependent on IRF3.

**Figure 2.**
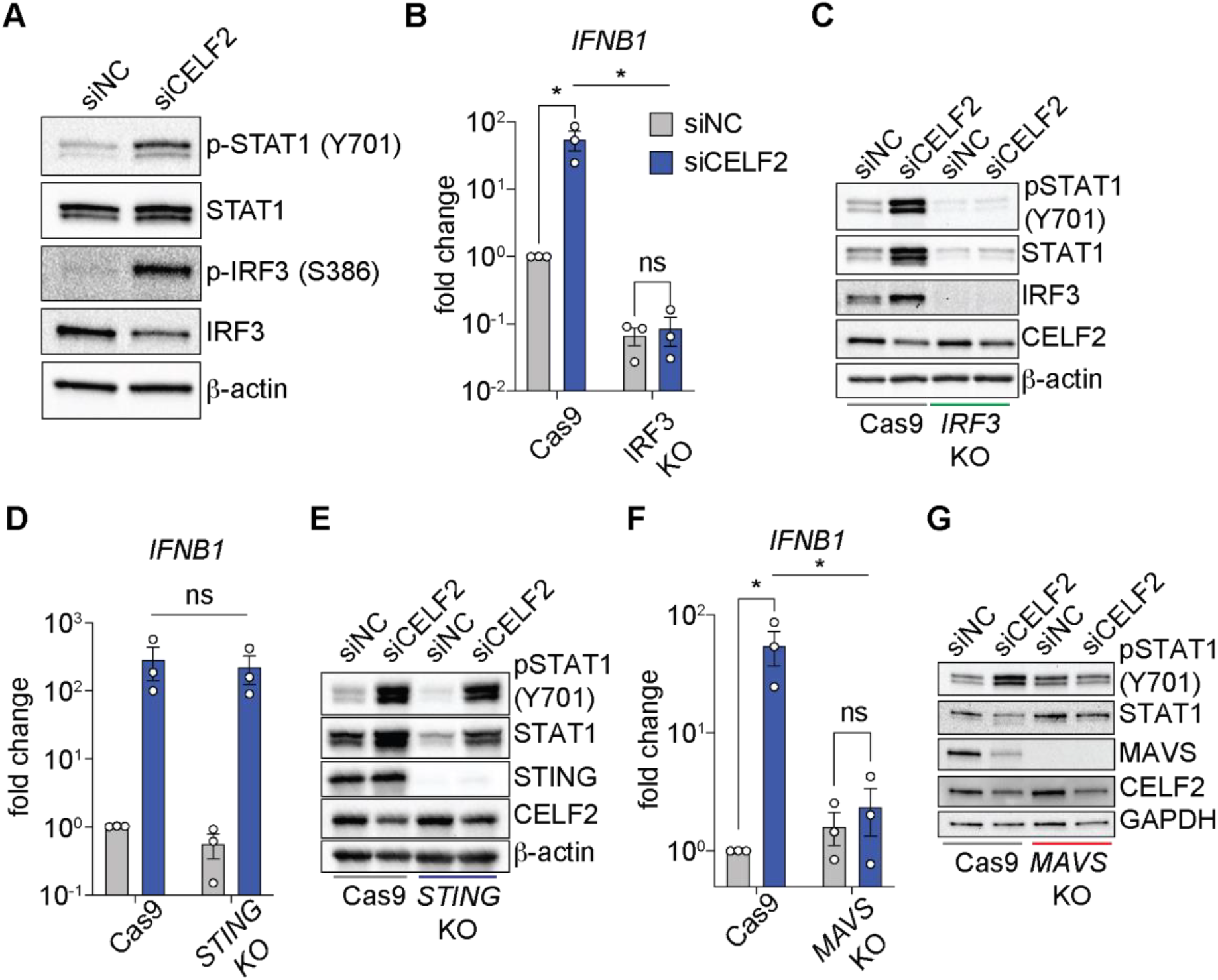
Spontaneous type I IFN and ISG expression following CELF2-depletion is RLR-dependent. A. Immunoblot analysis of STAT1 and IRF3 phosphorylation in CELF2-depleted cells. Representative of n=2 experiments. B-C. RT-qPCR of *IFNB1* expression and representative immunoblot analysis in IRF3 KO non-targeting control or CELF2-depleted cells. D-G. RT-qPCR of *IFNB1* expression and representative immunoblot analysis in STING (TMEM173) (D-E) or MAVS (F-G) KO control or CELF2-depleted cells. Data in all panels except A are n=3 independent experiments. Each symbol represents an individual biological replicate. Graphs display mean ± SEM. All data was analyzed by two-way ANOVA; ns represents p > 0.05, and * represents p < 0.05.

As both RNA and DNA sensing pathways are upstream of IRF3, we investigated the upstream adaptors and sensors that activate IRF3 in the absence of CELF2. We first generated U-937 cells deficient in the adaptor proteins STING and MAVS, required for DNA and RNA sensing through cGAS and RLRs, respectively. CELF2 depletion in STING KO cells did not abrogate *IFNB1* induction or STAT1 phosphorylation relative to non-targeting cells (Fig. 2D-E). However, deletion of MAVS resulted in a complete loss of *IFNB1* transcription and STAT1 phosphorylation (Fig. 2F-G). Furthermore, we observed decreased MAVS expression in CELF2-depleted cells. MAVS is ubiquitinated and degraded during RLR signaling(*40*), suggesting that RLR signaling is active following the depletion of CELF2. Taken together, these data suggest that loss of CELF2 in macrophages activates the RNA-RLR-MAVS and not the DNA-cGAS-STING sensing pathway.

### Loss of CELF2 results in the production of immunostimulatory RNA which is sensed by RIG-I

We next tested if RNA derived from CELF2-depleted cells could induce interferons in naïve cells. We isolated RNA from CELF2-depleted and control cells and transfected this RNA into ISRE-GFP reporter A549 cells, which fluoresce green upon the transcription of IFNs(*41*). As a control, transfecting ISRE-GFP reporter cells with poly U/UC RNA, which activates RIG-I (RIG-I ligand)(*6*), induced robust GFP expression by flow cytometry compared to non-transfected cells or cells treated with transfection reagent alone, indicating that these cells robustly respond to immunostimulatory RNA (Fig. S2A). RNA degradation by the RNase L pathway following viral infection can produce RLR ligands(*42*), therefore, we validated the integrity of the RNA isolated from CELF2-depleted and control cells by gel electrophoresis to ensure we were not transfecting fragmented RNA which could trigger RLR responses. We observed similar cellular RNA integrity from both treatment conditions (Fig. S2B). We then transfected RNA from CELF2-depleted and control macrophages into the ISRE-GFP reporter cells and observed a dose-dependent increase in the percent of GFP-positive cells upon transfection of RNA isolated from CELF2-depleted cells, but not from control cell-isolated RNA (Fig. 3A-B). These data suggest that loss of CELF2 results in the aberrant production of immunostimulatory RNAs which induce IFN.

**Figure 3.**
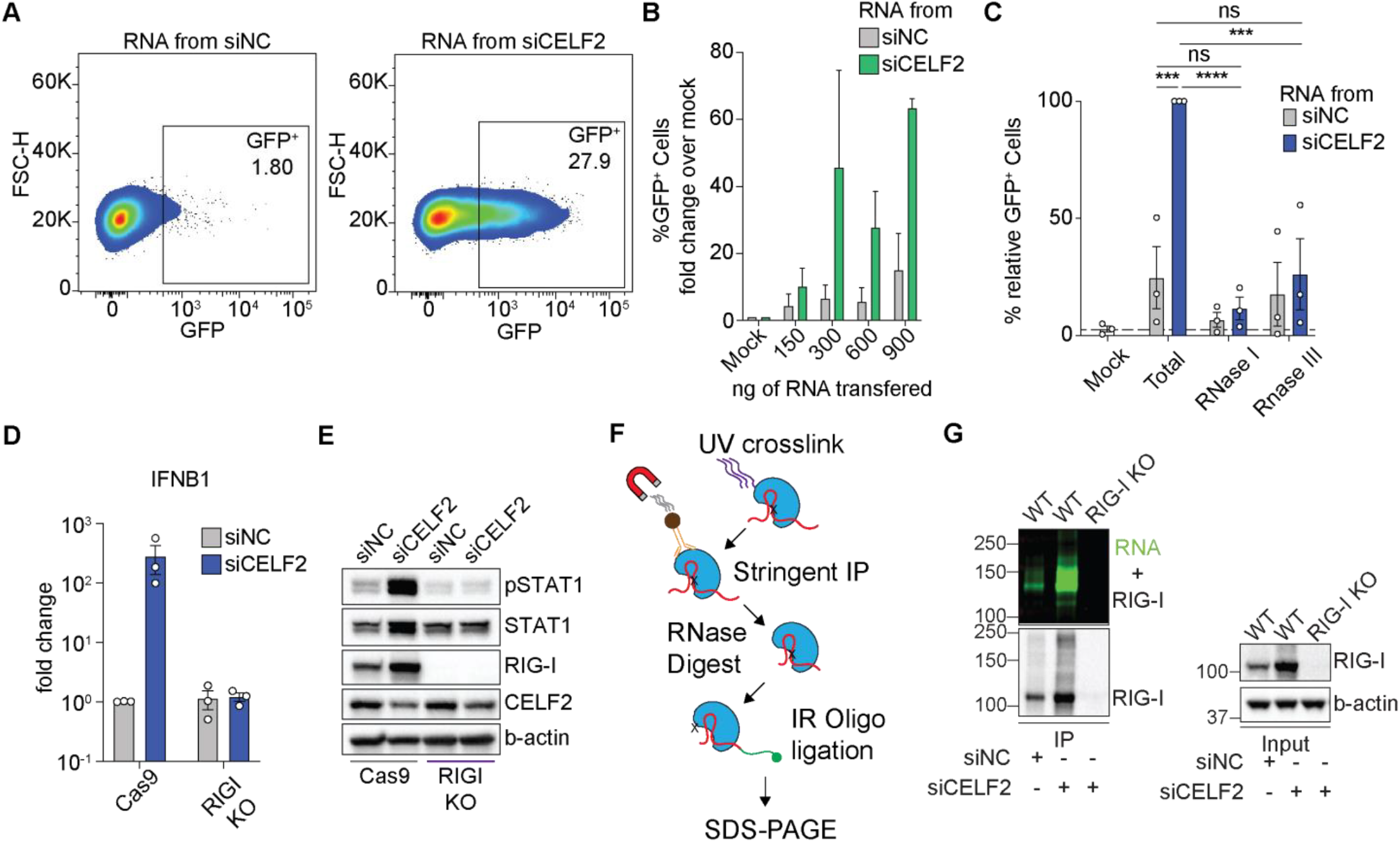
Loss of CELF2 results in the production of immunostimulatory RNA which is sensed by RIG-I. A. RNA transfer from CELF2-depleted and control cells into ISRE-GFP A549 cells. Flow plots are representative of 600 ng of transferred RNA and representative of 2 independent biological replicates. B. Dose-dependent expression of GFP in CELF2-depleted and control cells enumerated and representative of 2 independent biological replicates. C. Relative GFP expression in RNA transfer of total, RNase I, and RNase III treated RNA from CELF2-depleted and control cells. D-E. RT-qPCR of *IFNB1* expression (D) and representative immunoblot analysis (E) in RIG-I KO control or CELF2-depleted cells. F. Schematic of irCLIP experiment. G. irCLIP analysis of RIG-I:RNA complexes following CELF2-depletion. Data in all panels unless otherwise stated are n=3 independent biological replicates. Each symbol represents individual biological replicate. Graphs display mean ± SEM. B: two-way ANOVA, *** represents p ≤ 0.001 **** represents p ≤ 0.0001, ns represents p > 0.05. Immunoblots representative of n=3 independent biological replicates.

IFN can be induced following the sensing of specific RNA ligands, including dsRNA. To test if the loss of CELF2 results in the accumulation of endogenous immunostimulatory dsRNA, we digested RNA isolated from CELF2-depleted cells with RNase I or RNase III, which degrade single- and double-stranded RNA respectively. We confirmed the degradation of RNA via bioanalyzer; RNase I degraded almost all RNA, while RNase III did not (Fig. S2C). We then transfected either total RNA or RNase-treated RNA into ISRE-GFP reporter cells as previously described. As before, total RNA from CELF2-depleted cells significantly induced GFP expression compared to control cells, whereas RNase I treatment ablated the induction of GFP, likely due to the total degradation of isolated RNA (Fig. 3C). Importantly, we found that RNase III treatment of RNA reduced the induction of GFP to the levels of total RNA and RNase I-treated RNA (Fig. 3C). These data suggest that the loss of CELF2 results in the accumulation of endogenous immunostimulatory dsRNA.

Since IFN induction following CELF2 depletion relies on MAVS and IRF3, and dsRNA derived from CELF2-depleted cells can induce IFNs, we reasoned that immunostimulatory endogenous RNA generated in the absence of CELF2 must be sensed by either RIG-I or MDA5, or both. To test if RIG-I is required for IFN production following CELF2 depletion, we depleted CELF2 in RIG-I KO U-937 cells and measured the production of *IFNB1* and downstream activation of IFN signaling. We observed a complete loss of *IFNB1* induction and STAT1 phosphorylation following CELF2 depletion in RIG-I KO cells (Fig. 3D-E). On the other hand, siRNA depletion of MDA5 marginally reduced IRF3 phosphorylation after CELF2-depletion (Fig. S2D). Together, these data suggest that RIG-I is the primary sensor of immunostimulatory RNA produced following CELF2 depletion.

To test if an endogenous RNA ligand interacted with RIG-I, we performed infrared UV-C crosslinking immunoprecipitation (irCLIP), a method of visualizing RBP-RNA interactions via ligation of a fluorescent nucleic acid probe to RNA in complex with RBPs(*43, 44*). CELF2 was depleted in wild-type and RIG-I KO U-937s which were then exposed to UV to crosslink RNA to proteins. We then immunoprecipitated RIG-I and performed incomplete on-bead RNA digestion, and then ligated a fluorescent adapter which allowed for visualization of RIG-I:RNA complexes after SDS-PAGE electrophoresis (Fig. 3F). RNA-RBP complexes appear as a smear when visualized. We observed a distinct smear of RIG-I:RNA complexes following CELF2 depletion compared to that of control cells, indicating that RIG-I is bound to RNA in the absence of CELF2 (Fig. 3G). The fluorescent signal generated by RNA-RBP interaction was completely lost in RIG-I KO cells (Fig. 3G). Together these data indicate that the loss of CELF2 results in the accumulation of dsRNA which is sensed by RIG-I.

### Endogenous dsRNA ligands are enriched in CELF2-depleted cells

To identify the specific endogenous dsRNAs induced following the depletion of CELF2, we performed a dsRNA immunoprecipitation (dsRIP), wherein dsRNA was immunoprecipitated using the dsRNA-specific J2 antibody(*45*), isolated, and subjected to RNA-seq (Fig. 4A-C)(*46, 47*). Differential gene expression analysis of dsRIP input samples demonstrated a significant increase in ISGs as previously seen by long-read sequencing (Fig. 4A-B, 1B). Analysis of dsRNAs isolated from dsRIP revealed several transcripts significantly enriched in the CELF2-depleted macrophages compared to control cells (Fig. 4C). To test if any of the dsRNAs enriched in CELF2-depleted macrophages were contributing to the increased type I IFN signature observed in CELF2-depleted cells, we selected 12 candidate dsRNAs (6 coding and 6 non-coding) whose expression in the dsRIP input varied but were all enriched in the siCELF2 dsRIP (Fig. 4D). We then used siRNA to deplete 12 candidate RNAs in CELF2-depleted or control macrophages (Fig. 4E). Using *CXCL9, CXCL10* and *CXCL11* as representative ISGs, we found that single depletion of 3 mRNAs (*TM4SF20, SAPCD2*, and *VARS2*) and 2 non-coding RNAs (*LINC01031*, and *RNA5S17*) or all siRNAs combined resulted in a robust reduction in *CXCL10* and *CXCL11* expression in CELF2-depleted cells (Fig. 4E). These data suggest that specific dsRNAs enriched in CELF2-depleted macrophages contribute to the spontaneous IFN signature observed in CELF2-depleted cells.

**Figure 4.**
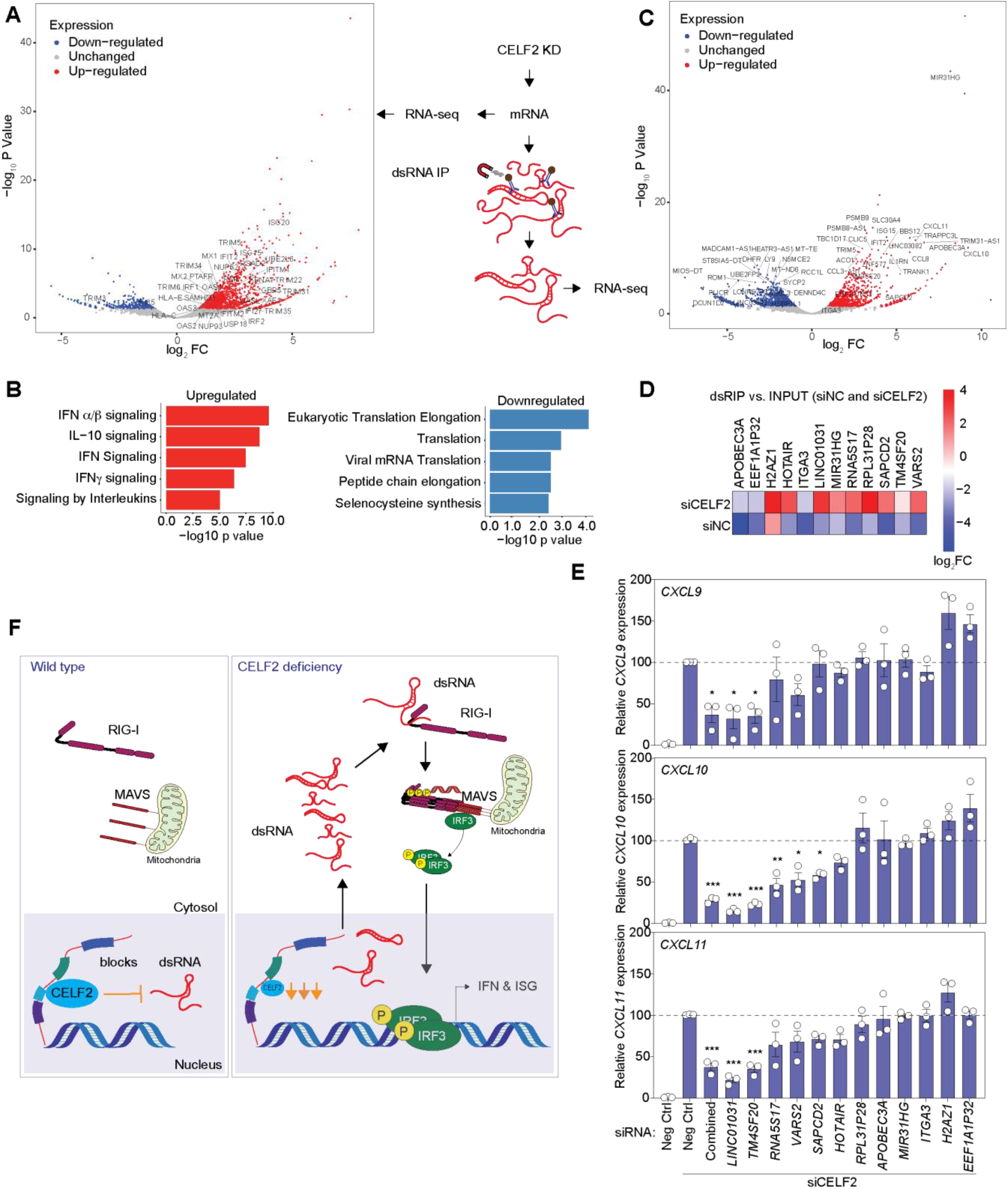
dsRNAs enriched in CELF2 depletion result in innate immune activation. A. Schematic of dsRNA IP and sequencing, right and volcano plot of ISGs upregulated following CELF2 KD (input), left B. Pathway analysis of significantly upregulated and downregulated genes from (A). C. dsRNA enrichment in CELF2-depleted macrophages compared to control macrophages following IP. For (A) and (C), red indicates dsRNAs significantly enriched in siCELF2; blue indicates dsRNAs significantly enriched in siNC, n=3. D. Heatmap of selected candidate expression from dsRIP in input. E. RT-qPCR analysis of *CXCL9, CXCL10* and *CXCL11* expression following knockdown of dsRIP candidates. F. A schematic showing that the splicing factor CELF2 suppresses endogenous dsRNA ligands which otherwise activate RIG-I signaling and produce an IFN signature.

## Discussion

In this study, we demonstrate that the loss of the RNA-binding protein CELF2 in macrophages leads to the spontaneous production of type I IFN. This process requires IRF3 and is driven by RIG-I-like receptor signaling, as deleting MAVS, but not STING, prevents the induction of interferon following CELF2 depletion. We show that CELF2 plays a role in suppressing the accumulation of dsRNA responsible for activating RIG-I, as RNase III treatment of RNA from CELF2-depleted cells significantly reduces its immunogenicity. Finally, we found that depleting several dsRNA ligands produced in the absence of CELF2 reduced ISG induction. Taken together, we identify CELF2 as a novel suppressor of endogenous dsRNA that can activate the RIG-I-like receptor pathway.

Cellular proteins that suppress endogenous RNA are critical in preventing IFN activation and the development of autoinflammatory diseases known as interferonopathies(*2*). While ADAR1, PNPase, and SKIV2L are known to edit or degrade dsRNA, regulators that suppress self-RNA biogenesis are unclear(*48*). CELF2, an mRNA splicing factor, has not been previously implicated as a suppressor of self-RNA ligands. CELF2 and other CELF family members bind to G/U-rich elements enriched in introns and 3’ untranslated regions. CELF2 has been shown to be critical for alternative splicing and alternative polyadenylation following T-cell receptor activation in T-cells(*27, 49, 50*) and downstream signaling of IL-10 in murine macrophages(*51*). CELF2 itself has not been directly implicated in the dysregulation of IFNs. The lack of reports indicating a role for CELF2 in IFN dysregulation could likely be attributed to the fact that loss of CELF2 in mice is embryonic lethal, and *de novo* mutations in humans are rare and result in severe neurological defects, making IFN phenotypes difficult to detect(*30, 34, 52, 53*). Interestingly, a study identified several single-nucleotide polymorphisms in CELF2 associated with acute respiratory distress syndrome, however this report did not investigate a mechanism of action(*54*).

Our observation that RNA in CELF2-depleted macrophages induces IFNs in a RIG-I-MAVS pathway-dependent manner strongly implicates CELF2 as a suppressor of endogenous RNA ligands. The direct role of CELF2 in immunity was corroborated by the finding that the RNA-binding ability of CELF2 was required for its suppression of IFNs, as overexpression of the RNA-binding mutant CELF2 failed to suppress spontaneous IFN production in CELF2-depleted cells. In previous studies, depletion of splicing factors like SF3B1(*55*), HNRNPM(*56*), and HNRNPC(*57*) has been implicated in the accumulation of dsRNA and IFN production. Some of these factors, such as HNRNPC and ADAR1, act as checkpoints to suppress MDA5 sensing and activation of IFNs(*57*). CELF2 and HNRNPC have several overlapping targets and coordinate splicing together in T cells, but whether CELF2 can replace HNRNPC and act as a checkpoint in coordination with ADAR1 is not known(*49, 58, 59*).Several dsRNAs of protein-coding genes were enriched following CELF2 knockdown, suggesting perturbations in mRNA splicing could be driving the IFN-stimulated phenotype observed in CELF2-depleted cells. This is distinct from ADAR1 suppression of MDA5 activation, which majorly edits non-coding regions. We down-selected candidate dsRNA ligands including coding and non-coding RNA species, which when depleted in CELF2-deficient cells, abrogated IFN-induced genes such as *CXCL9, CXCL10*, and *CXCL11*. These data implicate CELF2 as a central regulator responsible for suppressing the generation of dsRNA ligands and establish CELF2 along with other RBPs as a suppressor of aberrant IFN expression.

Within the context of an infection, CELF2 could protect the spliceosome complex from viral hijacking, in accordance with the guard hypothesis. This hypothesis proposes that hosts detect pathogens by monitoring disruptions in cellular function rather than through direct sensing of pathogens, a mechanism of defense employed by plants(*60, 61*). In mammalian cells, this was recently described within the context of HSV-1 infection, wherein the virus perturbs MORC3, leading to IFN activation(*62*). Future studies will investigate the possibility that pathogen disruption of CELF2 could effectively trigger IFNs, consistent with the guard hypothesis. Further, human genetic variations in *CELF2* are implicated in Alzheimer’s disease, autism, and impaired intellectual development(*63-66*), conditions which are often associated with dysregulation of IFNs. It would be interesting to investigate IFN responses in cell-specific CELF2 KOs or loss-of-function mutations within CNS disease models. Collectively, our study reveals a novel role for CELF2 in suppressing endogenous RNA that induces interferon, with implications in antiviral and autoimmunity.

## Materials and methods

### Cell lines, primary cells, cell culture conditions and treatments

U-937 monocytes were cultured in RPMI 1640 media supplemented with 10% FBS, 2mM glutamine, 100 U/ml penicillin and 100 mg/ml streptomycin and maintained at 37°C in 5% CO_2_. Non-targeted (H1), IRF3, STING, MAVS and RIG-I-deficient U-937 cells were generated by CRISPR-Cas9 genome editing by spinoculation lentiviral transduction and FLAG-GFP, FLAG-CELF2^WT^ and FLAG-CELF2^mut^ U-937 cells were generated by lentiviral transduction through spinoculation. Briefly, cells were incubated with specific lentiviruses packaged with pPAX and VSV-G, spun for 1 hour (h) at room temperature at 800 x g. Cells were then transferred and grown for 24h. Media was changed and transduced cells were incubated for another 24 h before enrichment using antibiotic selection. THP-1 monocytes were maintained in RPMI 1640 media supplemented with 10% FBS, 2mM glutamine, 100 U/ml penicillin, 100 mg/ml streptomycin, 1 mM sodium pyruvate, 10 mM HEPES and 0.05 mM, 2-mercaptoethanol and maintained at 37°C in 5% CO_2_. U-937 and THP-1 cells were differentiated for 48 h in their respective complete media containing 40 nM phorbol 12-myristate 13-acetate (PMA) followed by resting for 24 h in RPMI 1640 supplemented with 1% FBS. Primary monocytes were purchased from Bloodworks and differentiated in 50 ng/mL M-CSF in RPMI 1640 media supplemented with 10% FBS, 2mM glutamine, 100 U/ml penicillin and 100 mg/ml streptomycin for 7 days. Media was changed on day 3 with fresh M-CSF supplemented RPMI 1640 media. Huh7 human hepatoma cells, A549 human lung epithelial cells, HEK293T and derivative cell lines were cultured in DMEM supplemented with 10% FBS, 2mM Glutamine, 100 U/ml Penicillin and 100 mg/ml Streptomycin and maintained at 37°C in 5% CO_2_. Gene silencing was conducted using dicer-substrate interfering RNA (dsiRNA, IDT) or siGENOME SMARTpool siRNA (Dharmacon) specific to CELF2 or non-targeting control (Table S1). Transfections were carried out using 20 nM of dsiRNA or siRNA delivered intracellularly using TransIT-X2 (Mirus) (U-937, THP-1, Huh7, HEK293T) or Lipofectamine 2000 (Thermo Fisher) (primary monocytes) according to manufacturer’s guidelines.

### Plasmids and Oligonucleotides

CRISPR-Cas9 plasmids; pRRL-H1-PURO (non-targeting), pRRL-STING-PURO and pRRL-MAVS-PURO were a gift from Dr. Daniel Stetson (University of Washington)(*67*). pRRL-IRF3-PURO was generated in the lab as previously described(*68*) pRRL-RIGI-PURO was generated by cloning single-guide RNA (sgRNA) targeting RIG-I (5’-GTTCCTGTTGGAGCTCCAGG -3’) into empty pRRL-Cas9-PURO plasmids as previously described (*32, 67*) The ISRE reporter plasmid, pISRE-sfGFP, was a gift from Dr. Nicholas Heaton (Duke University) and has been previously described(*41*) pcDNA-FLAG-CELF2 (wild-type) was purchased from addgene. pCDNA-FLAG-CELF2^mut^ was generated using QuickChange Lightning Multi site-directed mutagenesis kit (Agilent) according to manufacturer’s guidelines. Primers for site directed mutagenesis can be found in Table S1. pLEX-FLAG-CELF2^WT^ and pLEX-FLAG-CELF2^mut^ were generated using infusion cloning (Takara), primers can be found in Table S1. Fluorescent oligo used for irCLIP can be found in Table S1.

### RNA extraction and quantification of gene expression

Total RNA was extracted using the NucleoSpin RNA extraction kit (Macherey-Nagel) or TRIzol reagent as indicated by manufacturer guidelines. cDNA synthesis was performed using the Prime Script RT (Takara Bio) according to the manufacturer guidelines. Relative quantification of mRNA was done by qPCR using the ViiA7 qPCR system with TaqMan reagents (Life Technologies) using *HPRT1* as a reference gene. Primers and probes used for qPCR assays in this study were acquired from IDT or Life Technologies as indicated in Table S1.

### RNA transfer experiments

RNA was isolated from CELF2-depleted or control cells as described. RNA was either digested with 1 U of RNase I/μg of total RNA or 1 U of RNase III/μg of total RNA of total RNA for 1h at 37°C or left untreated. RNase treated RNA was repurified using Phenol:Chloroform:Isoamyl alcohol extraction. RNA was transfected at indicated doses using Mirus TransIT X2 (Mirus) according to manufacturer guidelines.

### ISGF3 *Gaussia* luciferase and ISRE GFP reporter assays

5xISGF3 Huh7 cells were generated as previously described (*33*). Briefly, to generate the ISGF3 *Gaussia* Luciferase (Gluc) reporter construct, pTRIPZ-5xISGF3-BS-hGLuc-PEST, 5 tandem ISGF3 consensus sequences (5’-CGAAGAAATGAAACT-3’) were cloned with hGLuc-MODC-PEST into a pTRIPZ lentiviral plasmid. Lentivirus encoding the reporter was packaged and used to transduce human hepatoma Huh7 cells prior to single-cell cloning of reporter cells. To assess the presence of secreted IFN from dsiRNA transfected U-937 cells, cell supernatants from CELF2-depleted and non-targeting control treated cells were harvested 24 h post transfection and transferred onto 5xISGF3-GLuc Huh7 reporter cells. Reporter cells were then incubated at 37°C and 5% CO_2_ for 24 h prior to assessment of *Gaussia* luciferase secretion into the media. Sample supernatants were diluted 1:1 with *Gaussia* Luciferase glow assay substrate (Thermo Fisher Scientific) and luminescence was measured using a Synergy HTX (BioTek). To generate ISRE-GFP reporter cell lines, A549s were stably transduced with lentivirus pISRE-sfGFP and pools were transfected with RNA from CELF2-depleted or control cells as described and GFP expression was measured using a CANTO analyzer (BD). Flow cytometry data was analyzed using FlowJo (TreeStar).

### Western blot analysis

Whole cell lysates were prepared from cells using RIPA buffer (10 mM Tris-Cl (pH 8.0), 1 mM EDTA, 0.5 mM EGTA, 1% Triton X-100, 0.1% sodium deoxycholate, 0.1% SDS, 140 mM NaCl) supplemented with Halt protease and phosphatase inhibitor cocktail (Pierce). Protein quantification and normalization was done using the Quick-Start Bradford assay (Bio-Rad). 10-30 μg of total protein were resolved by SDS-PAGE and transferred to PVDF membranes (Bio-Rad). Primary antibody incubations were done overnight with antibodies diluted in 3% BSA in TBS-T (Tris-buffered saline/Tween 20), and species-specific HRP conjugated secondary antibodies. Chemiluminescent image acquisition was performed using a ChemiDoc XRS+ (Bio-Rad).

### Oxford Nanopore Technologies (ONT) MinION long-read RNA sequencing library prep and analysis

Total RNA was isolated as previously described. Total RNA was subject to polyA mRNA purification using oligo-dT Dynabead mRNA purification kit (ThermoFisher) according to manufacturer guidelines. 1 ng of polyA enriched RNA underwent PCR-cDNA barcoding using the PCR-cDNA barcoding kit (ONT) according to manufacturer guidelines. Briefly, RNA was reverse transcribed using VN primers and strand-switching primers provided in kit. cDNA was then amplified using barcoded cDNA primers provided in the kit and then subject to exonuclease I treatment. cDNA was purified using AMPure XP beads (Beckman Coulter) according to manufacturer guidelines and cDNA was checked for size and quality using the Agilent Bioanalyzer System (Agilent Technologies). The rapid adapter provided with kit was added to the barcoded cDNA. MinION R9.4.1 flowcell quality check, priming and loading was conducted according to manufacturer guidelines and sequencing allowed to progress for 72 h. Reads were base called using Guppy base caller (ONT). Base-called reads were trimmed and aligned (minimap2) to the GRCh38.p14 genome. Differential gene expression and alternative splicing analysis were conducted using FLAIR, as previously described(*26*).

### Infrared UV-C Crosslinking ImmunoPrecipitation (irCLIP)

irCLIP protocol was carried out as previously described(*43, 44*). Briefly, CELF2-depleted and control wild-type or RIG-I KO cells were UV-crosslinked at 150 μJ x 100/cm^2^ and then lysed. RIG-I was immunoprecipitated (IP) using polyclonal rabbit serum overnight at 4°C. The next day, samples were stringently washed, RNA was digested on bead using 25 ng/mL RNase A for 15 min at 30°C. RNA ends were then dephosphorylated and an IR-dye oligo (Table S1) was ligated onto the free ends of the RNA. RNA:protein complexes were then run on an SDS-PAGE gel and transferred to a nitrocellulose membrane. IR-RNA:protein complexes were imaged on a LI-COR Odyssey CLx (LI-COR). HRP-conjugated antibody imaging was performed as described.

### dsRNA immunoprecipitation and analysis

dsRIP protocol carried out as previously described(*46, 47*). Briefly, cells were lysed in dsRIP lysis buffer (100 mM NaCl, 50 mM Tris-HCl pH 7.4, 3mM MgCl_2_, 0.5% IGEPAL CA-630) prior to immunoprecipitation of dsRNA using the dsRNA specific monoclonal antibody, J2 (*45*) or isotype control antibody. 5% of the input was saved for total RNA control. RNA was isolated using TRIzol as previously described. RNA quality was assessed by bioanalyzer. RNA underwent ribosomal RNA depletion rather than mRNA selection to capture dsRNAs that may lack a poly-A tail. High-throughput sequencing was conducted using the Illumina NextSeq 500 sequencer at a read-depth of ∼60 million reads/sample.

We used dsRIP-RNA-seq to explore double-stranded RNA abundance in macrophages with CELF2 knockdown. RNA sequencing data was aligned to the human genome (hg38) with the Rsubread package. Count matrices were obtained with the feature Counts package. We normalized data using the trimmed mean of the M-values method using edgeR’s calcNormFactors function(*69*). Differential gene expression analysis was implemented using edgeR’s generalized linear model with quasi-likelihood F-tests. Gene annotations were obtained from biomaRt. Volcano plots were used to display differentially expressed genes (log2 fold change > 1.5, P < 0.05). For pathway analysis, we used clusterProfiler to perform Gene Ontology and KEGG pathway enrichment with a p-value cutoff of 0.05, using org.Hs.eg.db as a reference. Results were visualized using dotplots showing the top 10 enriched categories.

### Quantification and statistical analysis

Statistical analysis was performed using GraphPad Prism 9.0 (GraphPad software La Jolla, CA). Statistical significance was calculated as indicated for each experiment and across all experiments, p-values of < 0.05 were considered significant and are indicated by asterisks (*).

### Public data and reagent availability

Sequencing data are deposited under the GEO accession numbers GSE292983 and GSE293035. Data for Figs 1B, 4A, 4C, & 4D are in Supplementary Table S2. The reagents generated in this study are available upon request from the corresponding author under UW Materials Transfer Agreement.

## Supporting information

Reagents

Data for Figs 1B, 4A, 4C, & 4D

## Acknowledgements

We thank Daniel B. Stetson and Nicholas Heaton for sharing reagents. We thank Jennifer Hyde, Daniel Blanco-Melo, Tristan Jordan and Emmanuelle Genoyer for helpful discussions. This work was supported in part by the National Institutes of Health (R.S.: AI176442, J.R.S.: 2T32AI106677-6, O.C.K.: 5T32AI007509-25, N.S.G.: K99AI175483, M.G.: AI179722 and AI183793).

## Author Contributions

Investigation and Formal Analysis (J.R.S., N.S.G., N.E., G.M., K.S., J.S., C.M.P., O.C.K., M.G.); Conceptualization (J.R.S., R.S.), Writing (J.R.S., R.S., O.C.K.), Supervision (R.S.), Funding Acquisition (J.R.S., R.S., O.C.K., N.S.G. M.G.).

## Supplemental Figures

**Figure S1.**
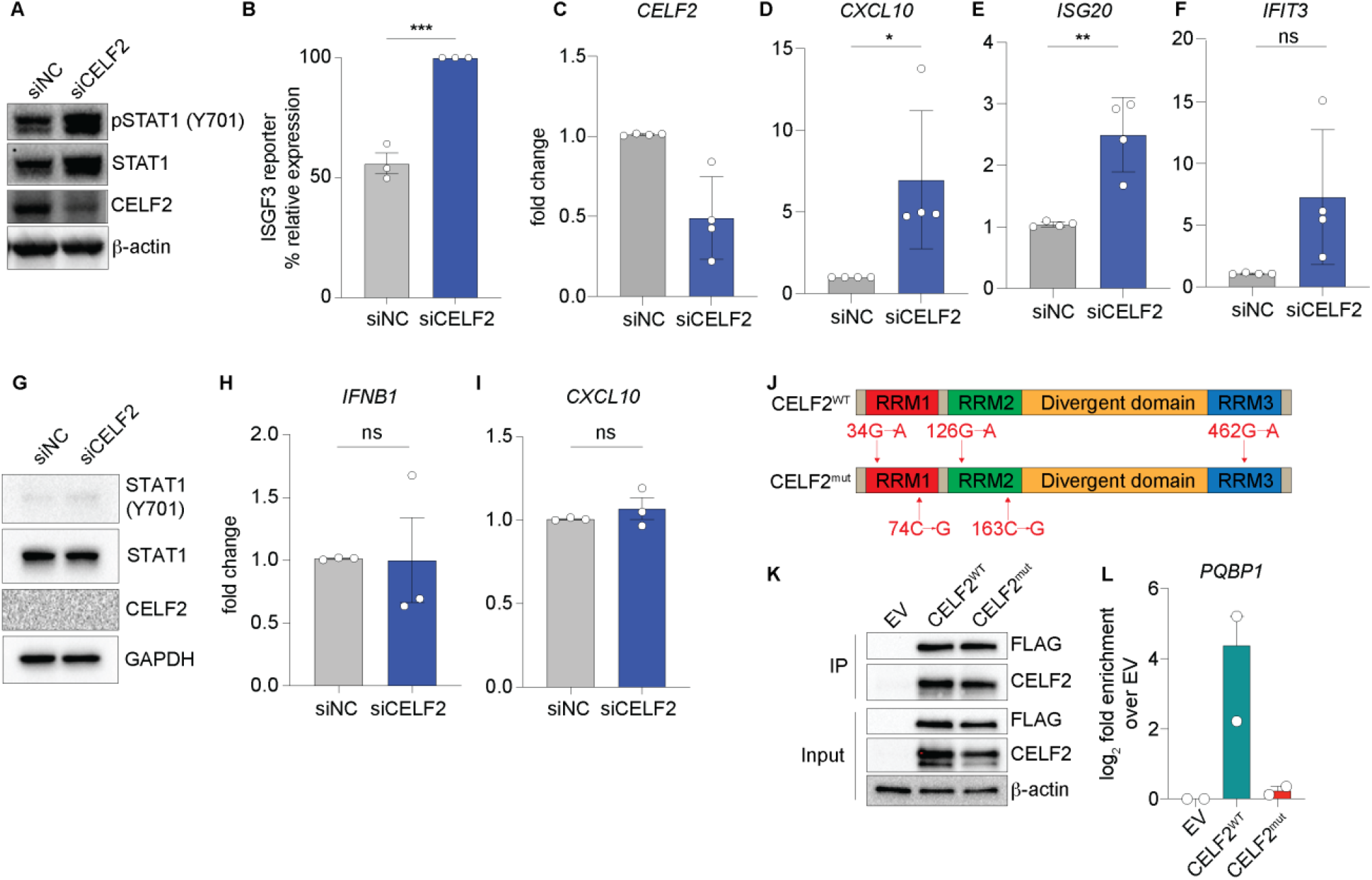
Type I IFN production following CELF2 depletion is specific to macrophages. A. Representative immunoblots of CELF2 depletion using pool of siRNA from Dharmacon. B. 5xISGF3 *Gaussia* luciferase reporter activity from CELF2-depleted U-937 using Dharmacon siRNA pools. C-F. RT-qPCR of *CELF2* (C), *CXCL10* (D), *ISG20* (E) and *IFIT3* (F) expression in CELF2-depleted THP-1 macrophages. n=4 independent replicates. G. Representative immunoblot analysis of CELF2 targeting in Huh7 cells. H-I. RT-qPCR of *IFNB1* (H) and *CXCL10* (I) expression in *CELF2*-targeted Huh7 hepatocytes. J. Schematic of CELF2 protein and amino acid mutations for CELF2 RNA-binding mutant. K. Representative immunoblot analysis of RNA immunoprecipitation (RIP) of FLAG-CELF2^WT^ and FLAG-CELF2 RNA-binding mutants (CELF2^mut^) with empty vector (EV control). L. RT-qPCR of RNA pulled down in CELF2 RIP for a representative gene. n=2. Data unless otherwise stated are n=3 independent biological replicates. (D-F, H-I) Students t-test; ** represents p ≤ 0.01, * represents p ≤ 0.05, ns represents p > 0.05.

**Figure S2.**
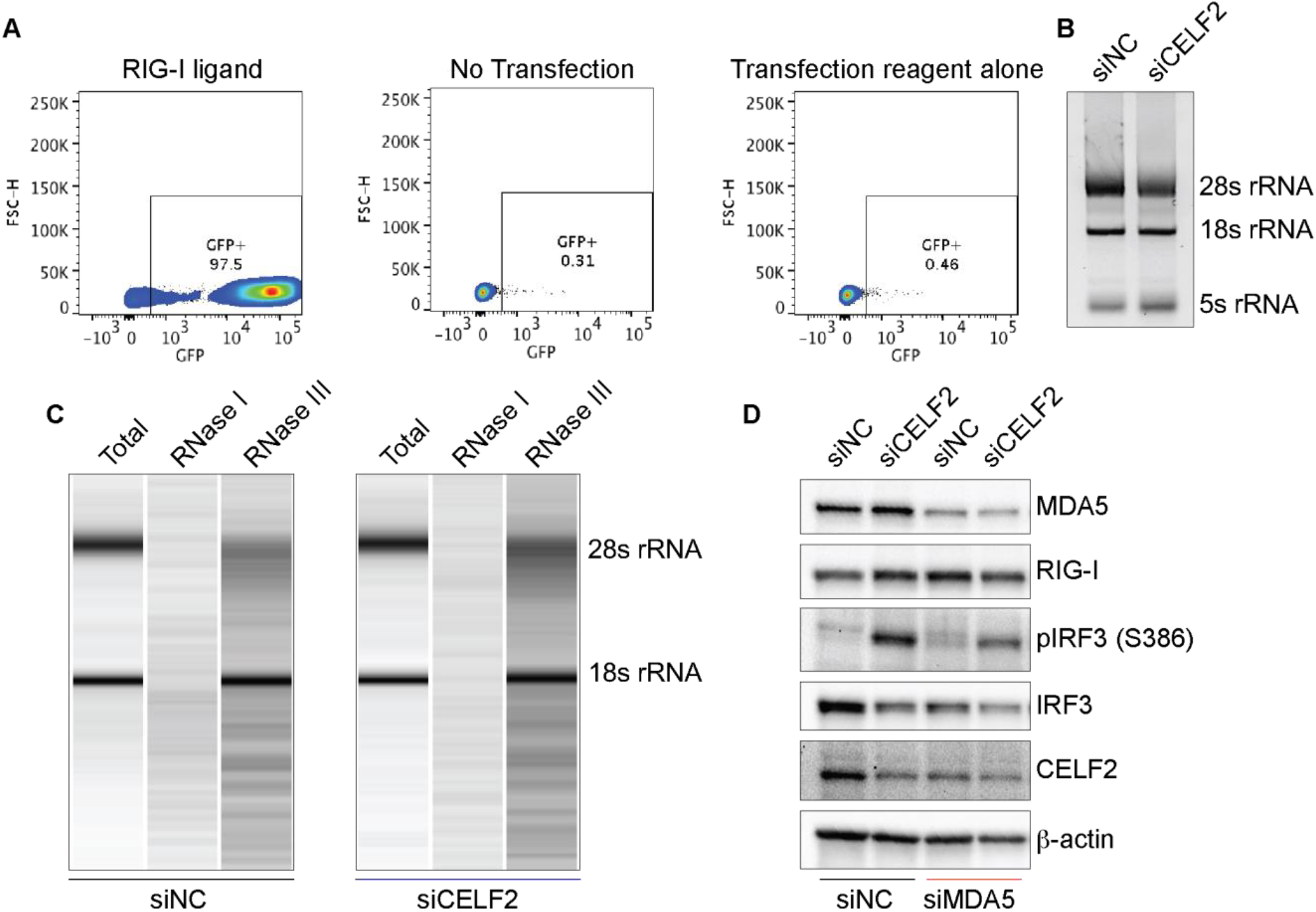
RNA controls in CELF2-depleted RNA transfer experiments. A. Representative flow plots of ISRE-GFP reporter using RIG-I Ligand and negative controls (no transfection or transfection reagent only). B. Representative gel analysis of RNA integrity of RNA from CELF2-depleted and control cells. 28s, 18s and 5s abundance was used to indicate RNA integrity. C. Bioanalyzer analysis of RNA degradation following treatment with RNase I and RNase III. D. Representative immunoblot analysis of IRF3 activation following MDA5 and CELF2 depletion.

